# Delta-catenin is required for cell proliferation in virus positive Merkel cell carcinoma cell lines but not in human fibroblasts

**DOI:** 10.1101/2025.03.12.642815

**Authors:** Joselyn Landazuri Vinueza, Nicholas J. H. Salisbury, Kristine N. Dye, Ann Roman, Denise A. Galloway

## Abstract

Merkel cell carcinoma (MCC) is a highly aggressive neuroendocrine skin cancer often driven by the integration of Merkel cell polyomavirus (MCPyV) into the host genome and the persistent expression of its viral oncoproteins, small tumor (ST) antigen and truncated large tumor (t-LT) antigen. While human fibroblasts support MCPyV replication, the cell of origin for MCC remains unknown. We hypothesized that MCPyV initially replicates in fibroblasts but, in rare cases, infects Merkel cell progenitors, contributing to MCC development. Using TurboID mass spectrometry, we identified δ-catenin as a novel ST interactor in fibroblasts. However, while ST binds δ-catenin in fibroblasts, this interaction is absent in virus-positive (VP)-MCC cell lines. Despite this, δ-catenin is essential for VP-MCC, but not for fibroblast, cell proliferation. We found that fibroblasts predominantly express δ-catenin isoform 1, whereas VP-MCC cells mainly express isoform 3. Overexpression of isoform 1 in VP-MCC failed to restore ST binding. δ-catenin promotes VP-MCC proliferation by regulating cell cycle gene expression through its interaction with Kaiso, a transcriptional repressor. Additionally, we found that LSD1 (KDM1A) regulates δ-catenin isoform 3 expression by modulating ESRP1, a δ-catenin splicing factor. Our findings reveal novel host factors involved in MCPyV infection and MCC tumorigenesis, suggesting that the host cell supporting viral replication and the MCC cell of origin may be distinct cell types.

**Importance:** Merkel cell polyomavirus (MCPyV), the only known human oncogenic polyomavirus, is the primary cause of Merkel cell carcinoma (MCC), a rare and aggressive type of skin cancer. MCC is driven by two viral proteins: small T (ST) and large T (LT). While the virus can replicate in skin fibroblasts, it is still unknown which type of skin cell becomes cancerous. We found that ST binds to a host protein, δ-catenin in fibroblasts, potentially playing a role in the virus lifecycle, but this interaction is missing in the cancer cells. Our study provides evidence that the cells in which the virus replicates and causes cancer are different.

## INTRODUCTION

Merkel cell carcinoma (MCC) is a rare and aggressive neuroendocrine skin tumor with two distinct etiologies: virus-negative (VN) and virus-positive (VP) MCC. VN-MCC tumors are caused by chronic ultraviolet (UV) exposure and are associated with high mutational burden frequently involving mutations in p53 and RB1 (1, 2). VP-MCC tumors are driven by the clonal integration of Merkel cell polyomavirus (MCPyV) genome into the host genome and the expression of the truncated large T (t-LT) and small T (ST) antigens, which are essential for tumorigenesis (3). Tumor-specific LT truncations inactivate LT’s DNA helicase function, disrupt viral replication while preserving the LXCXE motif, which is essential for binding to and inhibiting the tumor suppressor RB1 (4). In contrast, ST remains full length and forms a complex with L-Myc and EP400 that upregulates MDM2 and CK1α to activate MDM4 to suppress p53 activation (5). Furthermore, ST in complex with L-Myc and the EP400 complex, activates the expression of LSD1 (lysine-specific histone demethylase 1), also known as KDM1A, and other CoREST complex members including RCOR2 and INSM1. This complex mediates oncogenic transformation and tumorigenesis by repressing the expression of master regulators of the neuronal linage in MCC while promoting tumor cell proliferation (6, 7). In addition, several studies have shown that the expression of ST alone is sufficient to transform rodent fibroblast cells (8, 9) and human primary fibroblast (10) in vitro. Surprisingly, ST transformation functions are independent of the conserved interaction with protein phosphatase 2A (PP2A), seen with the oncogenic simian polyomavirus 40 (SV40) (8, 11). Furthermore, ST has shown oncogenic activity in mouse models (12, 13) indicating that ST is the dominant transforming viral protein.

The cell of origin of MCC remains unknown. MCC tumors were initially thought to arise from Merkel cells because they share expression of neuroendocrine markers. However, Merkel cells are highly differentiated and non-proliferative (14). As a results, other candidates have been suggested such as keratinocytes (HFK). The co-expression of ST and ATOH1, a Merkel cell linage transcription factor, is sufficient to initiate development of epidermis-derived MCC-like tumors in mice (15). Other candidates such as epidermal and dermal stem cells in addition to skin-derived precursors have been suggested (16) although none has been conclusively linked to MCPyV infection or development of MCC. Recently, MCC-like tumors were observed upon co-expression of MCPyV T antigens and ATOH1, in addition to the knockout of p53 in a keratin 5 (KRT5) promoter driven-mouse model (15). Interestingly, Merkel cells originate from basal epidermal progenitor cells that express both keratin 14 (KRT14) and through ATOH1-dependent transcriptional reprograming (17). However, whether these basal epidermal cells or their descendants, Merkel cells, are the cell of origin of MCC remains unclear.

So far, the only cell type that is permissive for the MCPyV lifecycle in vitro is human dermal fibroblasts (human foreskin fibroblasts or HFFs) (18). Typically, permissive cells support viral replication without undergoing transformation as observed in other polyomaviruses such as SV40 (19). This prompted us to investigate the cellular interactors of ST in HFFs using proximity labeling mass spectrometry (TurboID MS). Notably, since establishing an MCPyV infection model in cell culture remains technically challenging due to the difficulty of propagating MCPyV, we focused instead on expressing ST in HFFs. Among potential interactors, δ-catenin exhibited the strongest association with ST in HFFs. Additionally, δ-catenin has been shown to be required in the life cycle of human papillomavirus (HPV) (20). However, the role of δ-catenin in the life cycle of MCPyV and oncogenesis was unknown.

δ-catenin, also known as p120 catenin and encoded by the CTNND1 gene, is part of the catenin protein family, which also includes β-catenin. Human δ-catenin isoforms, referred to as isoform 1 through 4, differ from each other by the start codon used. Additional isoforms are derived from combinations with alternatively used exons A and B near the C-terminal end of the open reading frame and also with exon C in the middle of the open reading frame (21, 22). The relative expression of each isoform is different depending on the tissue and cell type. Moreover, the specific role of each isoform, especially isoforms 2 and 4, is poorly understood. Each isoform shares the central armadillo domain, which is essential for binding with cadherins on the cell membrane (22, 23) or transcription factors such as Kaiso in the nucleus (24). Therefore, the functional differences between the isoforms may be attributed to the N-terminus. δ-catenin isoform 1 being 101 amino acid residues longer at the N-terminus compared to isoform 3, potentially enables unique protein–protein interactions specific to isoform 1 (21, 22, 25). While δ-catenin isoform 1 has been reported to share regulatory mechanisms with β-catenin, the regulatory mechanisms governing δ-catenin isoform 3 are unclear. δ-catenin isoform 1 harbors GSK3β and CK1α phosphorylation sites within its unique N-terminal domain, which are absent in isoform 3. These sites facilitate β-TrCP recognition and subsequent degradation (26). It was unknown which δ-catenin isoforms are expressed in HFFs and MCC cell lines as well as how these isoforms are regulated.

While δ-catenin isoform 1 is highly expressed in mesenchymal cells, isoform 3 predominates in epithelial cells (25). During epithelial-mesenchymal transition (EMT) and cancer progression, transcription factors, such as Slug and Snail, promote isoform 1 expression (27), while Epithelial Splicing Regulating Proteins, ESRP1 and ESRP2, favor isoform 3 in epithelial tissues (28, 29). δ-catenin binds to cadherin at the plasma membrane through its armadillo domain supporting cell-cell adhesion (23). Loss, downregulation of or changes in δ-catenin phosphorylation prevents its binding with E-cadherin contributing to cell dissemination and is linked to metastasis and poor prognosis in several human cancers (23, 30). δ-catenin has also been found in the nucleus as it has two putative nuclear localization signals that interact with transcription factors including Kaiso. By directly interacting with Kaiso, δ-catenin indirectly activates the Wnt/β-catenin pathway (24). Kaiso is a BTB/POZ protein that has a bi-modal DNA-binding activity: it binds methylated CpG dinucleotides or sequence-specific consensus Kaiso binding sites within target gene promoters (31, 32) such as the Wnt/β-catenin genes Wnt11, Cyclin D1 and Matrilysin 7, to promote cell proliferation and migration (33–35). However, the role of δ-catenin in MCC was unknown. Interestingly, proteomic analysis identified δ-catenin among the most downregulated proteins following LSD1 inhibition, further suggesting its regulation by LSD1 (6). However, the mechanisms still are not understood. We hypothesized that while MCPyV initially infects HFFs for production of MCPyV virions, in rare instances, MCPyV may get into Merkel Cell progenitors contributing to the development of MCC. This prompted us to investigate the cellular interactors of ST in HFFs. We found δ-catenin exhibited strong binding to ST in HFFs while this interaction was absent in VP-MCC cells. Nonetheless, δ-catenin is essential for VP-MCC proliferation by regulating expression of genes involved in cell proliferation such as Cyclin B1 via interaction with Kaiso. Furthermore, we found that LSD1 regulates the expression of the δ-catenin isoform 3 by modulating ESRP1 expression, thereby promoting cell proliferation in VP-MCC.

## RESULTS

### Proximity labeling mass spectrometry identifies δ-catenin as a potential interactor of ST in fibroblasts

Human dermal fibroblasts are the primary host cell type productively infected by MCPyV (18), prompting us to identify ST interactors in human foreskin fibroblasts (HFFs) using proximity labeling mass spectrometry (36). HFFs were transduced with lentiviral vectors encoding TurboID (TID) biotin ligase fused to the N-terminus of ST (TID-ST). Since ST localizes to the nucleus without a known nuclear localization signal, likely through binding its interactors (9), we tested if TID fusion affected its localization. Subcellular fractionations confirmed that, like ST, TID-ST remains nuclear, suggesting that it retains its ST function and interactome (Fig. 1A). HFFs expressing TID or TID-ST were labeled with biotin, and biotinylated proteins were purified using streptavidin beads for mass spectrometry. Of 635 detected proteins, 121 were enriched in HFFs expressing TID-ST, including known ST interactors EP400 and HSP90 (5), validating this approach (Fig. 1B, blue dots). Among these, seven candidates were selected for further investigation for their roles in oncogenic and tumor-suppressive pathways: AIP (37), CACYBP (38), δ-catenin (23, 30), ING3 (39), TNK1(40), YAP, and AMOT (41) (Fig. 1B, red dots).

**Figure 1:**
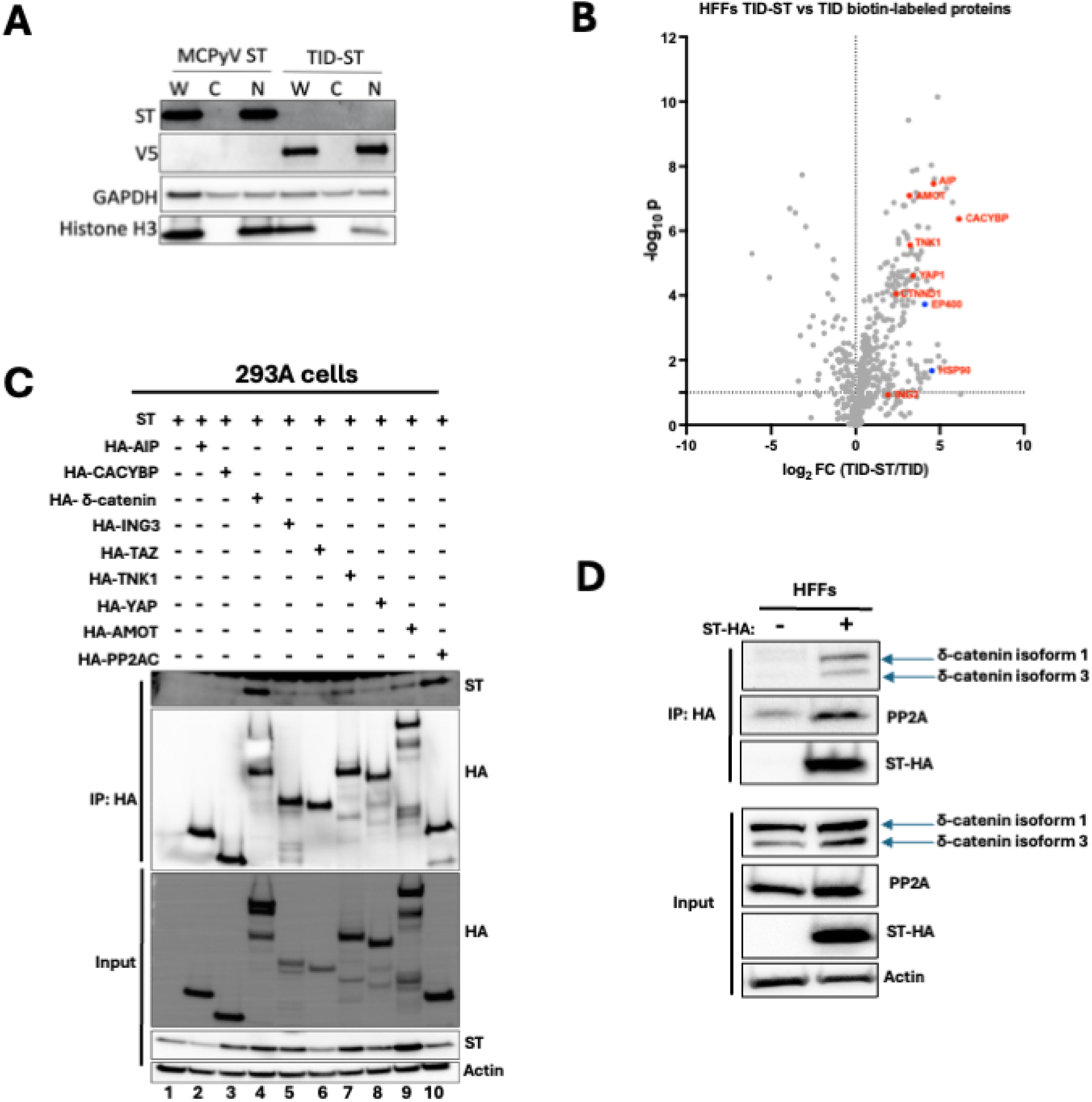
δ-catenin is a novel interactor of MCPyV ST in human foreskin fibroblast (HFFs). **A)** Subcellular localization of ST or TID-ST transduced into HFFs was detected with Ab5 antibody. TID-ST was detected with V5 antibody. Cytoplasmic and nuclear fractions were confirmed by GAPDH and histone H3 markers, respectively. W: whole lysate. C: cytoplasmic fraction. N: Nuclear fraction. **B)** Volcano plot of proteins identified by mass spectrometry enriched in HFFs TID-ST vs TID samples. Known interactors of MCPyV ST are highlighted in blue. Putative MCPyV ST interactors are highlighted in red. **C)** Western blots of HEK293A cell lysates transfected with ST and HA-AIP, HA-CACYBP, HA-δ-catenin, HA-ING3, HA-TNK1, HA-YAP, HA-AMOT or HA-TAZ. Lysates were immunoprecipitated with HA antibody and blotted for MCPyV ST. **D)** Western blots of HFFs cell lysates expressing either empty vector (-) or MCPyV ST-HA (+). Lysates were immunoprecipitated with HA antibody and blotted for δ-catenin.

To validate these interactions, each candidate was HA-tagged and co-transfected with ST in 293A cells followed by immunoprecipitation with an anti-HA antibody. ST co-immunoprecipitated with PP2AC, our positive control (8) (Fig. 1C, lane 10), and most of the putative interactors (Fig. 1C, lanes 2-9), though AIP and CACYBP showed no binding (lane 2,3), and YAP, TAZ and AMOT bound weakly (lanes 6, 8, 9). The strongest interaction was observed with δ-catenin (lane 4), leading us to focus our attention on δ-catenin.

To confirm the ST-δ-catenin interaction in HFFs, cells were transduced with a lentiviral vector encoding HA-tagged ST (ST-HA) or red fluorescent protein (RFP) as a negative control, followed by anti-HA immunoprecipitation and immunoblotting for endogenous δ-catenin. Consistent with the proximity labeling experiment, endogenous δ-catenin co-immunoprecipitated with ST-HA, but not RFP (Fig. 1D).

We then mapped the domain of δ-catenin interacting with ST. HA-δ-catenin isoforms 1-4 (Fig. S1A) were co-transfected with ST, followed by immunoprecipitation using an anti-HA antibody. ST bound strongly to isoform 1 (Fig. S1B, lane 3), weakly to isoforms 2 and 3 (lanes 4 and 5), and not to isoform 4 (lane 6) suggesting ST binds the N-terminal domain. Deletions mutants of isoform 1 (Fig. S1C) further defined the binding region. Isoform 1 served as a positive control, whereas isoform 4 served as a negative control (Fig. S1D, lanes 2 and 3). The Δ55-102 mutant had reduced binding (lane 4), while Δ103-324 mutant completely lost binding to ST (lane 5) suggesting that ST binds within the regulatory domain of δ-catenin. The Δ55–324 and the Δ325-968 mutants also failed to bind to ST, indicating that neither the coiled-coil domain alone (lane 6) nor the coiled-coil and regulatory domain together (lane 7) were sufficient for binding.

Given that the regulatory domain is heavily phosphorylated (42), we tested if phosphorylation was required for interacting with ST by generating a phospho-dead isoform 1 mutant (ΔP). Co-transfection of ST with ΔP in 293A cells showed that phosphorylation was not required for ST binding (Fig. S1E, lane 4).

Together, these findings suggest that ST binds δ-catenin within the regulatory domain, likely within the region spanning amino acids 102 to 324, and this interaction occurs independently of its phosphorylation. While the coiled-coil domain is not sufficient for binding, it may enhance the binding mediated by the regulatory region.

### MCPyV ST does not bind to δ-catenin in VP-MCC cells

Given the cell-type specificity of δ-catenin isoforms, we investigated their expression in MCC cells. Mesenchymal cells primarily express isoform 1, whereas epithelial cells predominantly express isoform 3 (25, 27). To determine which isoforms are expressed in MCC, the expression of δ-catenin was analyzed in a panel of 12 MCC cell lines, including variant VN-MCC and classical VN- and VP-MCC cell lines. Classical MCC cells grow as spheroids in suspension and express neuroendocrine markers (CHGA, SYP, ENO2) and Merkel cell markers (KRT20, ATOH1, SOX2) while variant MCC cells adhere in culture and show low or no expression of these markers (43)(Fig. S2A, S2B). δ-catenin isoform expression varies across these cell lines. Human foreskin keratinocytes (HFK) and classical VP-MCC cells predominantly expressed δ-catenin isoform 3, whereas HFFs and variant MCC cells expressed primary δ-catenin isoform 1. Most classical VN-MCC cells did not express δ-catenin (Fig. 2A). qRT-PCR analysis using isoform-specific primers (Fig. 2B), as expected, showed that HFFs and variant MCC cells expressed mostly isoform 1 mRNA (Fig. 2C, top) while HFK and classical VP-MCC cells predominantly expressed isoform 3 mRNA (Fig. 2C, bottom). Most classical VN-MCC cells show minimal δ-catenin mRNA, suggesting silencing of CTNND1 gene in these cells.

**Figure 2:**
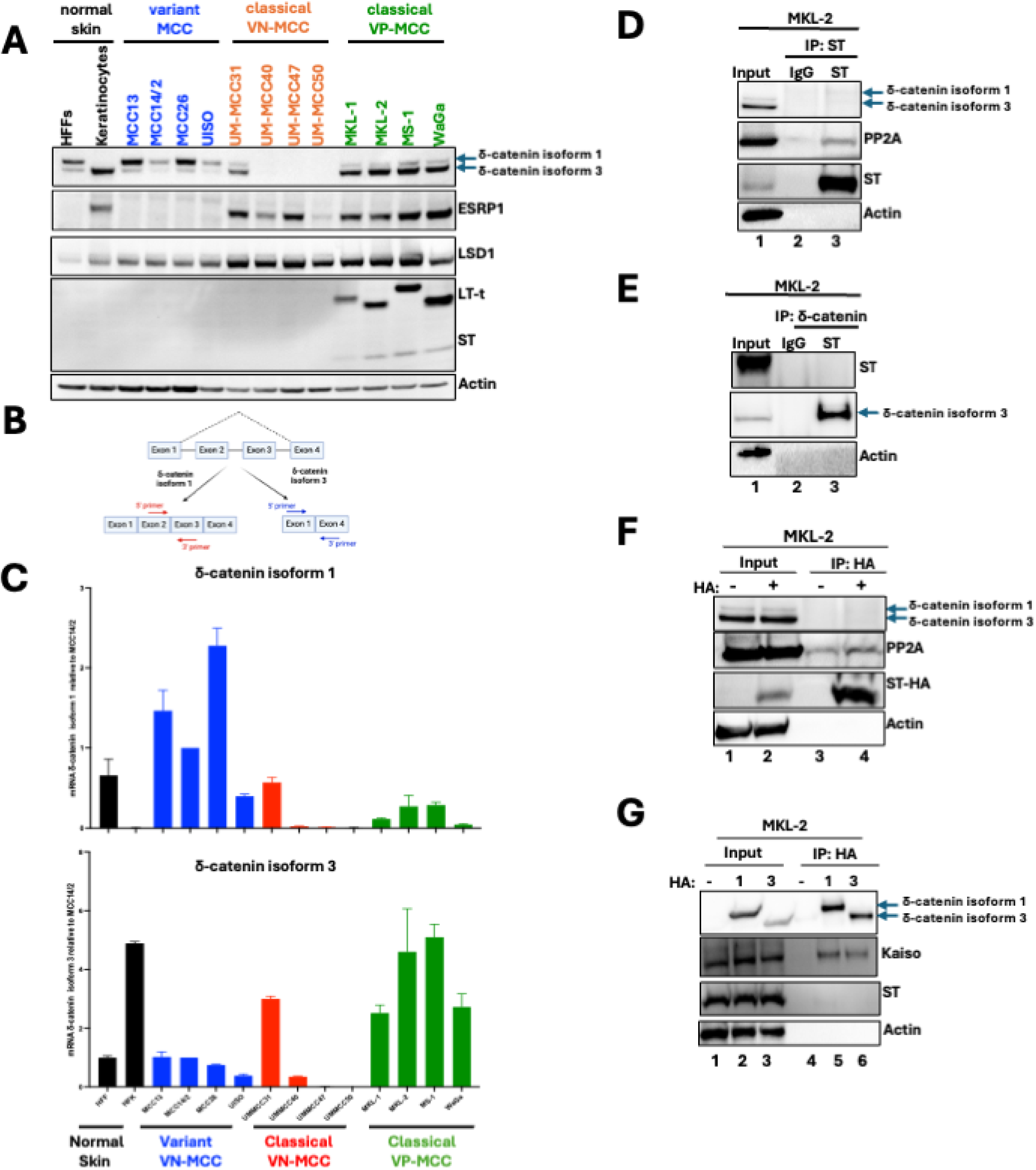
Failure of MCPyV ST binding to δ-catenin isoform 3 in VP-MCC Cells. **A)** Lysates from normal, variant, and classical virus negative and virus positive cells were immunoblotted for δ-catenin isoforms, ESRP1, LSD1 and MCPyV T antigens. **B)** Schematic representation of primers to detect δ-catenin isoform 1 and 3. Primers were designed to target exon-exon junctions specific to each isoform. **C)** qRT-PCR analyses of relative expression of δ-catenin isoform 1 and 3 in all 14 cell lines normalized to 36B4 expression. Data are shown as mean of n=3 ±SD. **D and E)** Western blots of MKL-2 lysates and anti-ST and anti-δ-catenin co-immunoprecipitations, respectively. **F)** Western blots of MKL-2 lysates overexpressing ST-HA and anti-HA co-immunoprecipitations after 6 days of doxycycline treatment. **G)** Western blots of MKL-2 lysates overexpressing HA-δ-catenin isoform 1 and 3 and anti-HA co-immunoprecipitations after 6 days of doxycycline treatment.

To test ST-δ-catenin interaction in VP-MCC cells, co-immunoprecipitation assays were performed using Ab5 antibody to pull down endogenous ST from MKL-2 whole-cell lysates. δ-catenin did not co-immunoprecipitate with Ab5 (Fig. 2D, lane 3), despite successful co-immunoprecipitation of PP2AC, a known ST interactor (lane 3). To confirm these results, a reciprocal co-immunoprecipitation was performed using an anti-δ-catenin antibody. Again, no interaction between endogenous ST and δ-catenin was observed in MKL-2 cells (Fig. 2E, lane 3).

To rule out the possibility that the lack of interaction was due to the antibodies used, ST was HA-tagged and overexpressed in MKL-2 cells using a doxycycline-inducible lentiviral vector followed by anti-HA immunoprecipitations. Again, endogenous δ-catenin did not co-immunoprecipitate with HA-ST in MKL-2 cells (Fig. 2F, lane 4). Given that VP-MCC cells express low δ-catenin isoform 1 levels compared to HFFs, (Fig. 1D), HA-tagged δ-catenin isoforms 1 and 3 were overexpressed in MKL-2 cells using a doxycycline-inducible lentiviral vector, and each of the isoforms was immunoprecipitated with an anti-HA antibody. ST still failed to bind either isoform (Fig. 2G, lanes 5 and 6) despite successful co-immunoprecipitation of Kaiso, a known interactor of δ-catenin (24) (lane 5 and 6). These findings suggest that ST and δ-catenin interaction requires more than δ-catenin isoform 1 being 101 amino acid residues longer at the N-terminus compared to isoform 3.

### δ-catenin is required for cell proliferation in VP-MCC cells

Although ST and δ-catenin do not interact in VP-MCC cells, we hypothesized that δ-catenin might still be necessary for cell proliferation, as it has been shown to promote cell proliferation in other cancers (33, 44, 45). To test this, MKL-1 and MKL-2 cells were transduced with two doxycycline-inducible lentiviral vectors encoding shRNA targeting δ-catenin, (Fig. 3A and 3B), and cell proliferation was assessed using trypan blue exclusion assays. δ-catenin knockdown significantly reduced cell proliferation in both MKL-1 and MKL-2 cells (Fig. 3C and 3D). This effect was specific to VP-MCC cells, as δ-catenin knockdown (Fig. 3E and 3F) did not affect proliferation in MCC14/2, a variant MCC cell line, or in HFFs (Fig. 3G and 3H). Notably, MCC14/2 cells and HFFs predominantly express δ-catenin isoform 1, whereas MKL-1 and 2 cells primarily express isoform 3. Since t-LT and ST are required for cell proliferation, we investigated whether δ-catenin knockdown affected their expression. However, δ-catenin knockdown had little to no effect on t-LT and ST in MKL-1 and 2 (Fig. 3A and 3B). Our findings indicate that δ-catenin is essential for cell proliferation in VP-MCC cells but not in HFFs and variant MCC and is completely dispensable in 3/4 VN-MCC cell lines.

**Figure 3:**
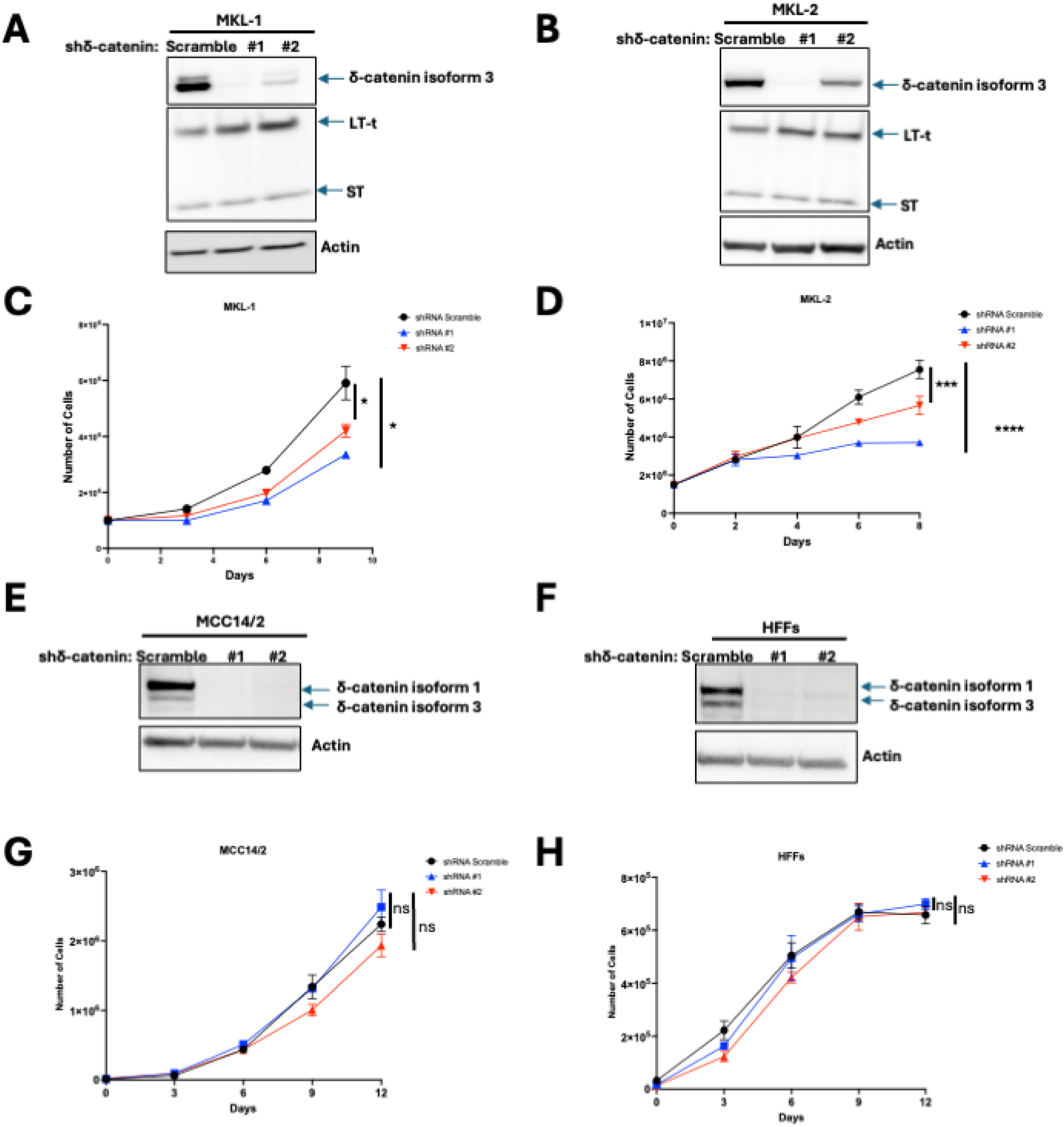
δ-catenin is required for cell proliferation in VP-MCC cells but not in MCC14/2 and HFFs cells. **A) and B)** Knock down of δ-catenin in MKL-1 and MKL-2. Western blots of cell lysates harvested after 8 days doxycycline treatment. **C) and D)** Growth curves of MKL-1 and MKL-2 knock down δ-catenin following doxycycline treatment. Data are shown as mean of n=3 ± SD; 2-way ANOVA, * P<0.05, ***P<0.001, ****P<0.0001. **E and F)** Knock down of δ-catenin in MCC14/2 and and HFFs. Western blots of cell lysates harvested after 8 days doxycycline treatment. **G and H)** Growth curves of MCC14/2 and HFFs knock down δ-catenin following doxycycline treatment. Data are shown as mean of n=3 ± SD; 2-way ANOVA, ns: no significant

### δ-catenin regulates genes involve in cell proliferation in MKL-2 cells

δ-catenin promotes cell proliferation by interacting with Kaiso, a transcriptional repressor of the Wnt/β-catenin pathway, thereby inhibiting its repressive activity (24, 34, 35). Consistent with this, endogenous Kaiso was successfully co-immunoprecipitated by δ-catenin isoform 3 in MKL-2 cells (Fig. 2G, lane 6). Kaiso regulates Wnt/β-catenin target genes, including Wnt11, Cyclin D1, and Matrilysin 7, in various cancers (33–35) but these were absent in classical VN and VP-MCC cell lines, while variant MCC cells expressed only Cyclin D1 (Data not shown). To identify δ-catenin target genes involved in proliferation, bulk RNA sequencing was performed following δ-catenin knockdown in MKL-2 cells. As expected, downregulation of cell cycle genes, including Cyclin B1 (46), Cyclin B3 (47), Cyclin A2 (48) was observed (Fig. 4A). Gene set enrichment analysis using the REACTOME collection showed that silencing δ-catenin in MKL-2 cells was associated with the downregulation of gene signatures related to cell proliferation (Fig. 4B).

**Figure 4.**
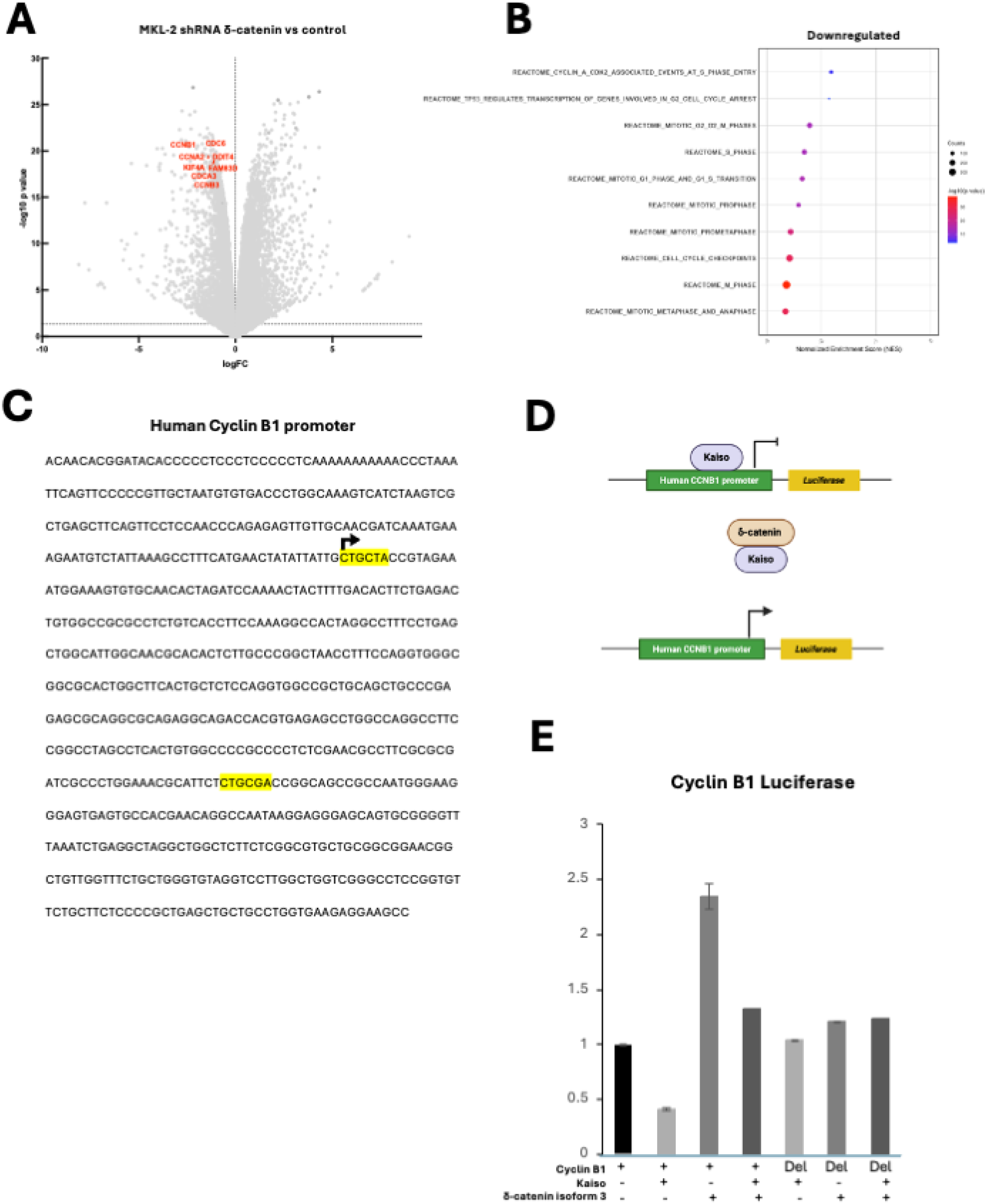
δ-catenin regulates expression of genes involve in cell cycle. **A)** Volcano plot of differentially expressed genes in MKL-2 knock down δ-catenin vs MKL-2 control cells from bulk RNA sequencing. **B)** Dot plot representing the changes in gene set enrichment analysis results illustrating processes associated with loss of δ-catenin in MKL-2. **C)** Nucleotide sequence of the human cyclin B1 promoter region. Consensus Kaiso binding sites are highlighted in yellow. Arrow indicates the transcription initiation site. **D)** Schematic representation of Kaiso and δ-catenin transcription regulation. **E)** Human CCNB1 promoter luciferase reporter in 293A cells co-transfected with Kaiso and/or δ-catenin isoform 3. Luciferase activity was normalized to CMV-driven renilla activity. Relative luciferase activity was determined using Cyclin B1 alone. Data are shown as mean of n=3 ± SD.

To determine if these genes are Kaiso target genes, we analyzed their promoters for Kaiso binding sites (KBS). Cyclin B1, B3, A2, and CDC6 contain KBS within their promoter regions (Fig. 4C and Fig. S3), suggesting potential regulation by Kaiso. As a proof of principle, we focused on Cyclin B1, a key regulator of the G2-to-mitosis transition (46). A luciferase reporter plasmid driven by the Cyclin B1 promoter was co-transfected into 293A cells along with Kaiso and/or δ-catenin isoform 3 (Fig. 4D).

As predicted, Kaiso overexpression repressed Cyclin B1 luciferase activity, whereas δ-catenin isoform 3 overexpression enhanced Cyclin B1 luciferase activity. Co-transfection of both Kaiso and isoform 3 attenuated Kaiso-mediated repression. To further confirm Cyclin B1 as a Kaiso target, we generated a luciferase reporter under the control of a Cyclin B1 promoter lacking KBS. Co-transfection of Kaiso, δ-catenin isoform 3, or both had negligible effects on luciferase activity (Fig. 4D), confirming that Kaiso binding is required for transcriptional regulation. Of note, we were unable to achieve efficient Kaiso knockdown in MKL-2 cells to assess its impact on cell viability and expression of cell cycle genes such as Cyclin B1 (data not shown).

Together, these findings suggest that δ-catenin promotes proliferation, at least in part, by alleviating Kaiso-mediated repression of Cyclin B1. The presence of KBS in the promoters of Cyclin B3 and A2 further suggests that these genes may also be potential Kaiso targets.

### LSD1 regulates the expression of δ-catenin isoforms through ESRP1

To investigate how δ-catenin isoforms are regulated, we examined the impact of LSD1 inhibition, as it has been shown that LSD1 inhibition alters δ-catenin expression (6). Treating MKL-2 cells with an LSD1-GSK inhibitor reduced cell viability in a dose-dependent manner (Fig. 5A). To confirm specificity, we analyzed PROM1 expression, a known LSD1 target, by western blot (6). As expected, LSD1 inhibition upregulated PROM1, validating LSD1 inhibition. Notably, LSD1 inhibition shifted δ-catenin expression from isoform 3 to isoform 1 (Fig. 5B). To confirm that this switch was specific to LSD1 inhibition, MKL-2 cells were transduced with LSD1-targeting shRNAs. Depletion of LSD1 resulted in a switch from δ-catenin isoform 3 to isoform 1 expression (Fig. 5C).

**Figure 5:**
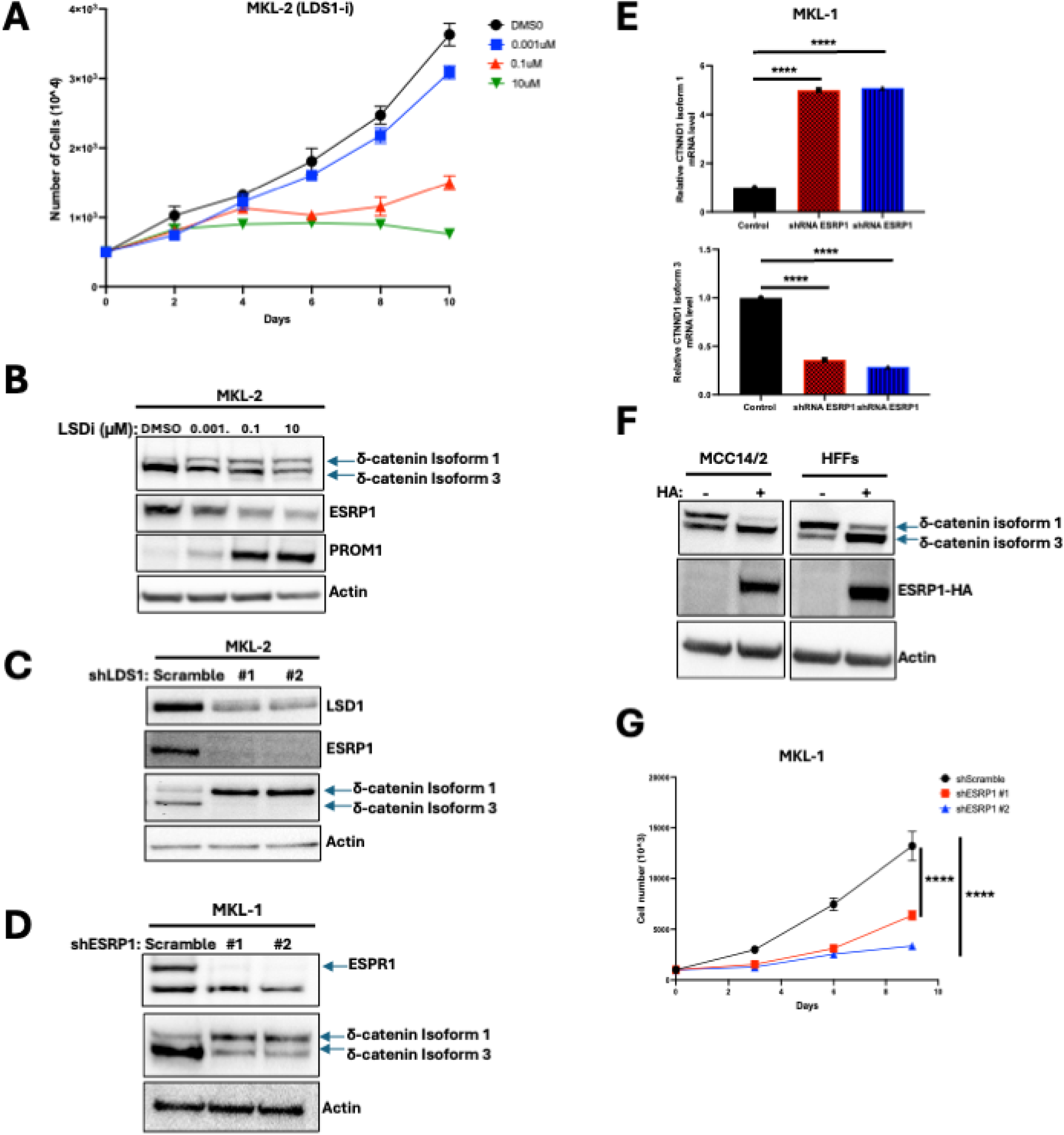
LSD1 regulates the expression of δ-catenin isoform 3 via ESRP1 in VP-MCC cells. **A)** Growth curve of MKL-2 after treatment with GSK-LSD1 inhibitor (LSD1-i). Data are shown as mean of n=3 ± SD. **B)** Western blots of δ-catenin, ESRP1 and PROM1 after 10 days treatment with LSD1-i in MKL-2 cells. **C)** Western blot of LSD1, ESRP1 and δ-catenin after 10 days knock down of LSD1 in MKL-2**. D)** Western blots of ESRP1 and δ-catenin after 10 days knock down of ESRP1 in MKL-1. **E)** qRT-PCR analyses of relative expression of δ-catenin isoform 1 and 3 after 10 days knock down of ESRP1 in MKL-1. Relative expression was normalized to 36B4. Data are shown as mean of n=3 ± SD; 2-way ANOVA, ****P<0.0001. **F)** Western blots of overexpressing HA tagged ESRP1 in MCC14/2 and HFFs cells after 10 days of doxycycline treatment. **G)** Growth curve of MKL-1 knock down ESRP1. Data are shown as mean of n=3 ± SD; 2-way ANOVA, ****P<0.0001.

Since ESRP1 regulates δ-catenin isoform expression (29), we hypothesized that LSD1 modulates δ-catenin splicing through ESRP1. HFK and classical MCC cell lines, which mostly express isoform 3, express ESRP1 while HFFs and variant MCC cells lines, which mostly express isoform 1, lack ESRP1 (Fig. 2A). Supporting this, LSD1 inhibition and knockdown in MKL-2 cells each decreased ESRP1 expression, leading to a switch from isoform 3 to isoform 1 (Fig. 5B and 5C).

To confirm ESRP1’s role in δ-catenin splicing, MKL-1 cells were transduced with ESRP1-targeting shRNAs. Knockdown of ESRP1 resulted in a switch from δ-catenin isoform 3 to isoform 1 (Fig. 5D), as confirmed by qRT-PCR analysis (Fig. 5E). Conversely, overexpression of HA-tagged ESRP1 in MCC14/2 and HFF cells, which lack ESRP1 and predominantly express isoform 1, shifted δ-catenin splicing from isoform 1 to isoform 3 (Fig. 5F). Additionally, ESRP1 knockdown (Fig. 5D) reduced proliferation in MKL-1 cells (Fig. 5G). ST induces the expression of LSD1 (6, 7) and our findings also add that LSD1 regulates δ-catenin isoform expression in VP-MCC cells through ESRP1, which is both necessary and sufficient for δ-catenin isoform 3 expression (Fig. S4) to promote cell proliferation in VP-MCC cells.

## DISCUSSION

While HFFs have been identified as a cell type permissive for the MCPyV viral lifecycle (18), the MCC cell of origin has yet to be determined. Typically, permissive cells are not transformed by viruses. For example, the related polyomavirus SV40 does not cause tumors in its natural host, rhesus macaque. Indeed, SV40 infection of permissive cells does not result in transformation because the intracellular environment supports viral replication, egress, and eventual cell death (19). In contrast, SV40 infection of non-permissive cells, such as hamster and mouse cells, is abortive, eliminating viral replication functions, and driving cellular transformation (49, 50). Using HFFs, a permissive cell for MCPyV infection, we identified δ-catenin as an ST interactor (Fig. 1B, C). While δ-catenin bound ST in HFFs (Fig. 1D), this interaction was absent in VP-MCC cells (Fig. 3A). Interestingly, while δ-catenin was required for cell proliferation in VP-MCC cells (Fig. 3C, D), at least in part by regulating the expression of cell cycle genes through Kaiso (Fig. 4A, E), δ-catenin was dispensable in HFFs (Fig. 3G, H). Although previous studies have shown that δ-catenin isoform 3, but not isoform 1, binds Kaiso (51, 52), we found that both isoforms bound Kaiso, at least in VP-MCC cells (Fig. 2G).

Notably, while our proximity labeling spectrometry identified YAP and TAZ as ST interactors in HFFs (Fig. 1B, C), MCC cells do not express YAP or TAZ (53). Our findings plus those of others indicate that HFFs provide a physiologically relevant environment for ST and δ-catenin interaction as well as YAP and TAZ, which may play a role in the MCPyV lifecycle. Moreover, this interaction is absent in VP-MCC cells, suggesting that the MCC cell of origin is distinct from fibroblasts.

Several skin cell types have been proposed as the MCC cell of origin, including Merkel cells, keratinocytes, or epidermal progenitor cells. However, none of these has been conclusively linked to MCC development. MCC tumors not only express neuroendocrine markers but also epithelial markers (54, 55). Our study reveals that MCC tumors express the epithelial markers ESRP1 (29) and δ-catenin isoform 3 (25) both of which are absent in HFFs and variant MCC cell (Fig. 2A). Loss of ESRP1, as seen during LSD1 inhibition, causes a switch from δ-catenin isoform 3 to isoform 1 in VP-MCC cells (Fig. 5B, C), likely resulting in a loss of epithelial identify. LSD1 repression inhibits MCC differentiation, upregulates expression of pro-neuronal genes such as NEUROD1 and INSM1 and promotes cell death in VP-MCC cells (6, 7). Interestingly, δ-catenin isoform 1 has been implicated in neurogenesis (56–58) suggesting that δ-catenin isoform 1 may contribute to the up-regulation to these neuronal genes, as observed previously (6, 7). Of note, LSD1 inhibition in MCC14/2, a variant MCC cell that primarily express isoform 1, did not affect its expression (data not shown) suggesting that LSD1 specifically regulates the expression of isoform 3 in VP-MCC cells.

While the MCC cell of origin remains elusive, our data suggest that MCC may arise from a cell with epithelial characteristics. While HFKs express both ESRP1 and isoform 3 (Fig. 2A), they are not considered a likely MCC precursor because viral genes are not expressed following viral infection (18). Most likely, MCC originates from a basal epidermal progenitor cell, precursors of Merkel cells (17). These epidermal progenitor cells probably express ESRP1 and isoform 3, as MCC cells do. Given that MCPyV is unlikely to drive oncogenesis in a permissive cell such as HFF, one possibility is that, while MCPyV replicates in HFFs, it occasionally infects Merkel progenitor cells, a non-permissive cell, in which the transcriptional environment represents a dead-end for viral replication, favoring viral integration and cellular transformation. Our work supports the idea that two distinct cells are involved: HFFs supports the MCPyV lifecycle, while MCC may arise of epidermal progenitor cells.

## Acknowledgements

MKL-1 and MKL-2 cell lines were a gift from Drs. Yuan Chan and Patrick Moore and MS-1 cell line was a gift from Dr. Masahiro Shuda (University of Pittsburgh, Pittsburgh, Pennsylvania, USA). WaGa cell line was a gift from Dr. Jurgen Becker (University of Duisberg-Essen, Duisburg, Germany). UM-MCC31, UM-MCC40, UM-MCC-47, UM-MCC-50 cell lines were a gift from Drs. Monique E. Verhaegen and Andrzej A. Dlugosz (University of Michigan, Ann Harbor, USA). UISO cell line was a gift from Dr. Isaac Brownell (National Institute of Health, Bethesda, USA). Ab5 antibody, used to detect ST and LT, was a gift from Dr. James A. DeCaprio (Harvard University, Boston, USA). This research was funded by the National Institutes of Health/National Cancer Institute (grant #: R35 CA209979) to DAG and the Viral Pathogenies and Evolution Training Grant (grant#: T32 AI083203) to JLV and KD. Assistance with TurboID experiments was provided by Lisa Jones, Chen Wei Lin, and Phil Gafken and the Fred Hutch Proteomics & Metabolomics Resource, which is funded in part through NIH/NCI Cancer Center Support Grant P30 CA015704. This RNAseq data was supported by the Genomics & Bioinformatics Shared Resource, RRID:SCR_022606, of the Fred Hutch/University of Washington/Seattle Children’s Cancer Consortium (P30 CA015704) with assistance of Alyssa Dawson and Pritha Chanana.

## MATERIALS AND METHODS

### Cell lines

MCC13 (#10092302), MCC14/2 (#10092303) and MCC26 (#10092304) cells were from Sigma. Variant VN-MCC and VP-MCC cells were culture in RPM1 1640 HEPES (Thermo fisher) with 10% tetracycline-negative FBS (GeminiBio), 1X non-essential amino acids (Gibco) and 50 units/mL penicillin and 50 ug/mL streptomycin (Gibco). Classical VN-MCC cells were maintained in media containing 50% DMEM (Gibco), 30% Neurobasal medium (Gibco), 15% chick embryo extract (Thermo fisher), 1% N2 (Gibco), 2% B27 (Gibco), 117nM retinoic acid (Sigma), 20ng/ml bFGF (R&D), 20ng/ml IGF-1 (R&D), 1% non-essential amino acids (Gibco), 1% penicillin and streptomycin (Gibco) and 50uM 2-βmercaptoethanol (Sigma). 293A cells and 293T cells (Invitrogen) were maintained in DMEM (Gibco) with 10% FBS (Corning), 1x GlutaMAX (Gibco), 1X non-essential amino acids (Gibco), 50 units/mL penicillin and 50ug/mL streptomycin (Gibco). Fibroblasts were cultured in DMEM (Gibco) with 10% FBS (Corning), 1x GlutaMAX (Gibco), 1X non-essential amino acids (Gibco), 50 units/mL penicillin and 50ug/mL streptomycin (Gibco). Keratinocytes were cultured with keratinocyte growth medium 2 (PromoCell) per the manufacturer’s instructions. All cell lines were maintained at 37°C in 5% CO2.

### Plasmids

ST interactors, δ-catenin isoforms and δ-catenin mutants as well as ESRP1-HA were codon-optimized, synthesized as dsDNA fragments (IDT) and cloned into pcDNA3 or pTRIPZ using the In-fusion HD Cloning Kit (Takara).

### Transfections and lentiviral transductions

For transient transfections, 293A were plated in 10cm plates, at ∼80% confluence the next day, transfected with TransIT^®^-293 Mirus following the manufacturer’s instructions. Cells were harvested 36 hours post-transfection.

To generate lentiviral particles, 293T cells were transfected with psPAX2 and pMD2.G (gifts from Didier Trono, Addgene #12260, #12259) and lentiviral vectors (see Appendix, Table S1 or pTRIPZ (Adgene, #206981) expressing δ-catenin isoform 1-HA, isoform 3-HA, ST-HA or ESRP1-HA. After 36h, supernatants were collected, filtered through a 0.45μM filter, and supplemented with 6μg/mL polybrene. Target cells were incubated with lentiviral supernatants for 24h, followed by fresh medium for another 24h.

Transduced cells were then selected with 0.5μg/mL puromycin for 4 days before expansion in puromycin-free medium.

### Cell growth assays

Cells transduced with doxycycline-inducible lentiviral vectors were treated with 0.5ug/ml doxycycline, refreshed every 3 days. MKL-1 cells transduced with shRNA against ESPR1, and MKL-2 cells transduced with shRNA against LSD1 were refresh every 3 days. GSK-LSD1 (Sigma, #SML1072,) was reconstituted in DMSO and added to the media. At each time point, cells were treated with accutase (Stem Cell Technologies) for 10 minutes at room temperature and cell number was assessed by 0.2% trypan blue staining.

### Co-Immunoprecipitation, immunoblotting and antibodies

Cell were lysed in 1% NP-40 lysis buffer (50 mM Tris HCl pH 8.0, 150 mM NaCl, 1% NP-40) with protease and phosphatase inhibitors. Protein concentrations were determined using BCA protein assays (Thermo Fisher). For co-immunoprecipitations, 500ug of lysates were pre-cleared with 25ul protein A/G magnetic beads (Pierce) for 1h at 4°C with rotation, then incubated with 25ul anti-HA beads (Pierce) for 1h at 4°C. Beads were washed five times with 1%NP-40 buffer and bound . Bound proteins were eluted with 1x LDS sample buffer (Novex) plus 100 mM DTT at 70 °C for 10 min.

Twenty μg of cell lysate and IP elute were separated on 8 to 16% denaturing tris-glycine gels (Thermo Fisher), transferred to PVDF membranes (Bio-Rad), blocked in 3% BSA in TBS containing 0.1% Tween-20 (TBST) for 1 h at room temperature followed by incubation with primary antibodies overnight at 4°C. Blots were washed with TBST 3 times followed by HRP-conjugated secondary antibodies for 1 h at room temperature followed by washes with TBST 3 times. Blots were developed using Clarity Max Western ECL chemiluminescent substrate (Bio-Rad) and imaged on ChemiDoc (Bio-Rad).

For immunoblotting alone, cells were lysed in RIPA buffer (25nM Tris-HCL, 150mM NaCl, 1% NP-40 and 1% deoxycholate) supplemented with protease and phosphatase inhibitors and process as above. Antibodies used are listed in Appendix, Table S2.

### Microscopy

Bright-field microscopy images were taken at 10x magnification using the EVOS microscope.

### Luciferase assay

Human CCNB1 promoter (Appendix, Table S3) was cloned into the pGL4.11 luciferase vector (Promega). 2×10^4^ 293A cells in a 96 well plate were transfected with 1ug pGL4.1 CNNB1-luc, 1ug CMV renilla pGL4.75 (Promega) and 1ug pcDNA3 Kaiso and/or 1ug pcDNA3 δ-catenin isoform 3, with pcDNA3 empty vector added to equalized total DNA. Cells were lysed and analyzed using the Dual-Glo luciferase assay kit (Promega) on a Veritas microplate luminometer (Turner BioSystems).

### Subcellular fractionations

Subcellular fractionation was performed using a Thermo Fisher kit (#78840). Lysates were processed as described previously. Antibodies used are listed in Appendix, Table S2.

### RNA extraction and qRT-PCR

Total RNA was isolated using the RNeasy Mini Kit (Qiagen) per manufacturer’s protocol and quantified with a NanoDrop spectrometer. cDNA was synthesized using the SuperScript VILO cDNA Synthesis Kit (Thermo Fisher) according to the manufacturer’s instructions. qRT-PCR was performed using PowerUp SYBR Green Master Mix (Thermo Fisher) and StepOnePlus Real-Time PCR system (Applied Biosystems). Differential expression of δ-catenin isoform 1 and 3 was calculated using the 2−ΔΔCt method and normalized to housekeeping gene 36B4 expression. Primers sequences are listed in Appendix, table S4.

### Bulk RNA Sequencing

Total RNA was isolated using the RNeasy Mini Kit (Qiagen), with RNA integrity assessed by TapeStation (Agilent Technologies) and quantificatify using a Trinean DropSense96 spectrophotometer (Caliper Life Sciences). RNA-seq libraries were prepared with the TruSeq stranded mRNA kit (Illumina) using 500ng total RNA, and library size was validated using a TapeStation. Additional quality control, pooling of indexed libraries, and cluster optimization were performed using the Invitrogen Qubit® 2.0 Fluorometer (Life Technologies-Invitrogen). Libraries were pooled (9-plex) and clustered onto a P2 flow cell before sequencing on an Illumina NextSeq 2000 with a paired-end, 50 bp (PE50) strategy.

Image analysis and base calling were performed using Illumina’s Real Time Analysis v3.4.4 software, followed by demultiplexing and FASTQ generation with bcl2fastq Conversion Software v2.20. Reads were aligned to the GRCm38 reference using STAR v2.7.7a in two-pass mode, and gene quantification was performed with STAR’s quantMode using Gencode v38 annotations. Differential expression was analyzed using Bioconductor’s edgeR with a significance threshold of log₂FC ≥ 1 or ≤ −1 at 5% FDR. Gene Set Enrichment Analysis (GSEA) was performed on pre-ranked gene lists to identify enriched pathways.

### Proximity Labeling Mass Spectrometry

HFFs TID and ST-TID were cultured with biotin-free DMEM media (Gibco) supplemented with 10% FBS (Corning), 1x GlutaMAX (Gibco), 1X non-essential amino acids (Gibco), 50 units/mL penicillin and 50ug/mL streptomycin (Gibco). At 90% confluency, cells were incubated with DMEM containing 500 µM biotin for 30 minutes, then placed on ice, washed with ice-cold PBS, and collected. Cells were resuspended in ice-cold 1% NP-40 lysis buffer with benzonase (Sigma) and rotated for 30 minutes at room temperature. Protein concentrations were measured by BCA assay. Six mg of protein was incubated with 125 µL streptavidin magnetic beads (Pierce) that were pre-methylated using a reductive alkylation kit (Hampton Research HR2-434) for 2 hours at room temperature. Beads were washed twice with 1% NP-40 buffer, and proteins were eluted in 8 M urea/100 mM Tris (pH 8.5). Disulfide bonds were reduced with 5 mM TCEP for 15 minutes and alkylated with 10 mM 2-chloroacetamide for 30 minutes in the dark at room temperature. Samples were digested with 250 ng rLys-C at 37°C for 4 hours, diluted to 1 M urea with 100 mM Tris (pH 8.5), supplemented with 1 mM CaCl₂, and further digested overnight with 1.6 µg trypsin at 37°C. After acidification with 5% formic acid, peptides were desalted using ZipTips (Millipore), eluted with 70% acetonitrile/0.1% TFA, and dried in a Speedvac.

Peptides were resuspended in 2% acetonitrile in 0.1% formic acid and analyzed by LC/ESI MS/MS on a Thermo Scientific Easy1200 nLC coupled to a tribrid Orbitrap Eclipse with FAIMS Pro. In-line desalting was accomplished using a reversed-phase trap column (100 µm × 20 mm, Magic C18AQ, 5 µm, 200 Å; Michrom), followed by separation on a reversed-phase column (75 µm × 270 mm, ReproSil-Pur C18AQ, 3 µm, 120 Å; Dr. Maisch). A 90-minute gradient from 8% to 30% B (80% acetonitrile in 0.1% formic acid) at 300 nL/min was used. Data were acquired in a data-dependent mode with survey scans in the Orbitrap and MS/MS spectra in the linear ion trap (HCD activation) with dynamic exclusion for 60 seconds after one repeat.

Data analysis was performed using Proteome Discoverer 2.5 (Thermo Scientific). The data were searched against Uniprot Human (UP000005640_Human_052618.fasta) that included common contaminants (cRAPome, cRAP_012915_NP.fasta) and the ST sequence (PC_KD_1622.fasta). Searches were performed with settings for the proteolytic enzyme trypsin. Maximum missed cleavages were set to 2. The precursor ion tolerance was set to 10 ppm and the fragment ion tolerance was set to 0.6 Da.

Dynamic peptide modifications included oxidation (+15.995 Da on M). Dynamic modifications on the protein terminus included acetyl (+42.-11 Da on N-terminus) and static modification carbamidomethyl (+57.0215 on C). Minora was used for peak abundance and retention time alignment. SEQUEST HT was used for database searching. All search results were run through Percolator for scoring. Raw data was normalized at median value across samples. Missing data were imputed with half of global minimum values. P-values for pairwise comparisons were calculated by t-test.

